# REC8-cohesin, chromatin and transcription orchestrate meiotic recombination in the Arabidopsis genome

**DOI:** 10.1101/512400

**Authors:** Christophe Lambing, Andrew J. Tock, Kyuha Choi, Stephanie D. Topp, Pallas C. Kuo, Alexander R. Blackwell, Xiaohui Zhao, Kim Osman, James D. Higgins, F. Chris H. Franklin, Ian R. Henderson

## Abstract

During meiosis chromosomes undergo DNA double-strand breaks (DSBs) that can be repaired using a homolog to produce crossovers, which creates genetic diversity. Meiotic recombination occurs coincident with homolog pairing and polymerization of the meiotic axis and synaptonemal complex (SC). REC8-cohesin is required to connect chromosomes to the axis and to organize axis polymerization. However, control of REC8 loading along chromosomes, in relation to chromatin, transcription and recombination, is not yet fully understood. Therefore, we performed REC8 ChIP-seq in Arabidopsis, which revealed strong enrichment in centromeric heterochromatin. REC8 abundance correlates with suppression of meiotic DSBs and crossovers, despite axis loading of SPO11-1 in these regions. Loss of the heterochromatic marks H3K9me2 and non-CG DNA methylation in *kyp/suvh4 suvh5 suvh6* mutants causes remodeling of REC8 and gain of meiotic recombination locally in repeated sequences, although centromere cohesion is maintained. In the chromosome arms, REC8 is enriched within gene bodies, exons and GC-rich sequences, and anti-correlates with transcription. Highest REC8 occupancy occurred in facultatively silent, H3K27me3-modified genes. Using immunocytology we show that axis polycomplexes form in *rec8* mutants that recruit recombination foci with altered stoichiometry, leading to catastrophic non-homologous recombination. Therefore, REC8 plays a key role organizing meiotic chromosome architecture and promoting high-fidelity interhomolog recombination. Despite this pro-recombination role, local REC8 enrichment associates with DSB repression at the fine scale, which is consistent with the tethered-loop/axis model. Coincident with its organizational role during meiosis, REC8-cohesin occupancy along the chromosomes is shaped by multiple chromatin states and transcription.

## Introduction

Cohesin complexes form ~35-50 nm rings that can topologically embrace one or more DNA helices (Uhlmann 2016; Nasmyth and Haering 2009; Peters et al. 2008). Cohesin rings consist of paired structural maintenance of chromosomes (SMC) proteins that interact at hinge and ATPase head domains, with the head regions clamped by an α-kleisin (Gligoris and Löwe 2016). DNA can enter and exit cohesin rings at the subunit interfaces, and the rings undergo dynamic cycles of association and disassociation with chromosomes (Uhlmann 2016; Nasmyth and Haering 2009; Peters et al. 2008). Cohesin complexes regulate diverse nuclear processes, including chromosome condensation, segregation, gene expression and DNA replication, recombination and repair (Uhlmann 2016; Nasmyth and Haering 2009; Peters et al. 2008). Cohesin complexes containing the α-kleisin REC8 also play critical roles in controlling meiotic chromosome segregation and interhomolog recombination.

During meiosis eukaryotic genomes undergo DNA double-strand breaks (DSBs) generated by SPO11 topoisomerase-like complexes (Keeney et al. 1997; Baudat et al. 2013). Meiotic DSBs can enter an interhomolog repair pathway to produce reciprocal crossovers, which creates genetic diversity (Hunter 2015). Following meiotic S-phase, sister chromatids form co-aligned chromatin loops connected to axial element polymers via REC8-cohesin complexes (Zickler and Kleckner 1999). The meiotic axis includes HORMA domain proteins (e.g. ASY1) (Armstrong et al. 2002), and interacting partners (e.g. ASY3) (Ferdous et al. 2012), which promote interhomolog recombination. Although REC8-cohesin plays essential roles in establishing the tethered-loop/axis architecture, its role in regulation of meiotic recombination is complex. For example, budding yeast cohesin and axis components are required to additively promote meiotic DSB repair (Klein et al. 1999; Kim et al. 2010), whereas local Rec8 enrichment suppresses DSB formation in both budding and fission yeast (Panizza et al. 2011; Storlazzi et al. 2008; Nambiar and Smith 2018). As meiosis proceeds, HORMA proteins become depleted, as synaptonemal complex (SC) components are installed (e.g. ZYP1), until synapsis completes at pachytene (Ferdous et al. 2012; Lambing et al. 2015; Higgins et al. 2005). Hence, REC8-cohesin, axis and SC proteins play tightly integrated roles in establishing a chromosome structure that favors interhomolog repair during meiosis.

SPO11 and its accessory factors dynamically associate with the axis in fungi and mammals during DSB formation, and DNA repair occurs at axis-associated sites (Stanzione et al. 2016; Blat et al. 2002; Panizza et al. 2011; Baudat et al. 2013; Nambiar and Smith 2018; Pan et al. 2011; Sommermeyer et al. 2013; Acquaviva et al. 2013). Budding yeast DSB hotspots occur within nucleosome-free regions in the chromatin loops, in proximity to H3K4me3-modified nucleosomes at gene 5’-ends (Pan et al. 2011; Borde et al. 2009). Direct interactions occur between the Spp1 complex, which catalyzes and binds to H3K4me3, and the meiotic axis protein Mer2 (Sommermeyer et al. 2013; Acquaviva et al. 2013). This supports a model where loop DNA sequences are tethered to the axis via specific chromatin modifications during DSB formation and repair. Interactions occur between functionally related components in mammals, including H3K4me3, CXXC1, IHO1 and PRDM9 (Parvanov et al. 2017; Imai et al. 2017), and plant crossovers are positively correlated with gene density and H3K4me3 (Choi et al. 2013). Therefore, euchromatic marks likely play conserved roles in recruiting loop DNA to the axis during meiotic recombination. Transcription itself plays an important role in shaping cohesin accumulation in gene-rich regions in mitosis and meiosis, and can therefore also influence recombination (Lengronne et al. 2004; Bausch et al. 2007; Sun et al. 2015; Busslinger et al. 2017).

In contrast to euchromatin, meiotic recombination is typically suppressed in heterochromatin (Underwood et al. 2018; Choi et al. 2018; Ellermeier et al. 2010). Heterochromatin is repeat and transposon-dense, late-replicating, suppressed for RNA polymerase II transcription and densely modified with epigenetic marks including DNA and H3K9 methylation (Janssen et al. 2018). Heterochromatic marks including DNA and H3K9 methylation are sufficient to silence meiotic recombination hotspot activity in plants, fungi and mammals (Zamudio et al. 2015; Yelina et al. 2015; Maloisel and Rossignol 1998). Cohesin is also strongly enriched in the centromeres and pericentromeric heterochromatin of animals and fungi (Tanaka et al. 1999; Mizuguchi et al. 2014; Sun et al. 2015; Blat et al. 2002; Klein et al. 1999; Bernard et al. 2001; Watanabe and Nurse 1999). In fission yeast, centromeric cohesin enrichment requires heterochromatin protein Swi6 and H3K9 methylation (Bernard et al. 2001; Nonaka et al. 2002; Mizuguchi et al. 2014), implying a direct connection between heterochromatic epigenetic marks and cohesin recruitment. However, mouse centromeric cohesion is maintained during mitosis in *suv39h-1 suv39h-2* H3K9 methylation mutants, although local cohesin remodeling occurs on specific repeats (Koch et al. 2008; Guenatri et al. 2004). Therefore, the functional relationships between cohesin, chromatin state and transcription during meiosis, and the consequences for interhomolog recombination, remain incompletely understood.

Here we use a comprehensive array of genetic, genomic and immunocytological approaches to investigate the role of Arabidopsis REC8 in orchestrating meiotic chromosome architecture and recombination, and its functional interactions with chromatin and transcription. We show that *rec8* mutants undergo defective axis polycomplex formation, which associates with catastrophic non-homologous recombination. Using REC8 ChIP-seq we show strong enrichment in centromeric heterochromatin and within specific classes of RNA transposable elements. REC8 levels anti-correlate with meiotic DSBs (SPO11-1-oligos) and crossovers at both the chromosome and fine scales. To directly test the role of heterochromatin on cohesin loading, we performed REC8 ChIP-seq in *kryptonite (kyp/suvh4) suvh5 suvh6* triple mutants, which lose H3K9me2 and non-CG DNA methylation. We observed remodelling of cohesin and DSB landscapes in repeated sequences in *kyp suvh5 suvh6*, although centromere cohesion is maintained, meaning that Arabidopsis more closely resembles mice than fission yeast. Transcriptional changes in *kyp suvh5 suvh6* are associated with remodeling of REC8 occupancy and meiotic DSBs in both genes and transposable elements at the fine-scale. At the cytological scale, we show that *rec8* axis polycomplexes are able to recruit the recombination machinery, although with altered stoichiometry. The *rec8* polycomplexes undergo synapsis and ultimately cause non-homologous recombination. Hence, REC8-cohesin organizes meiotic chromosome architecture and high-fidelity homologous recombination, and is simultaneously influenced by both chromatin state and transcription.

## Results

### Complementation of *rec8* meiotic catastrophe via epitope tagging

To detect REC8 during meiosis we inserted 3×HA or 5×Myc epitopes into a genomic clone and transformed *rec8-3/+* heterozygotes. In wild type meiosis, chromatin matures from thin threads at leptotene, to thick paired axes at pachytene with condensed heterochromatin, until five bivalents connected via chiasmata are evident at metaphase I (Fig. 1A and Supplemental Fig. S1A). In contrast, no axis differentiation occurs in *rec8* chromatin, which is present as a diffuse mass during mid-prophase I although the proportion constituting heterochromatin is not significantly different (Mann-Whitney-Wilcoxon (MWW) test, *P*=0.55) (Fig. 1A, Supplemental Fig. S1A-S1B and Supplemental Table S1). The entangled chromatin mass in *rec8* proceeds to fragment at metaphase I, causing complete sterility (Fig. 1A) (Cai et al. 2003; Chelysheva et al. 2005). We observed that epitope-tagged *REC8* constructs complemented *rec8* meiotic phenotypes, including (i) axis formation during prophase I, (ii) the presence of five bivalents at metaphase I and (iii) chiasmata counts (Fig. 1A, Supplemental Fig. S2A and Supplemental Table S2).

**Figure 1.**
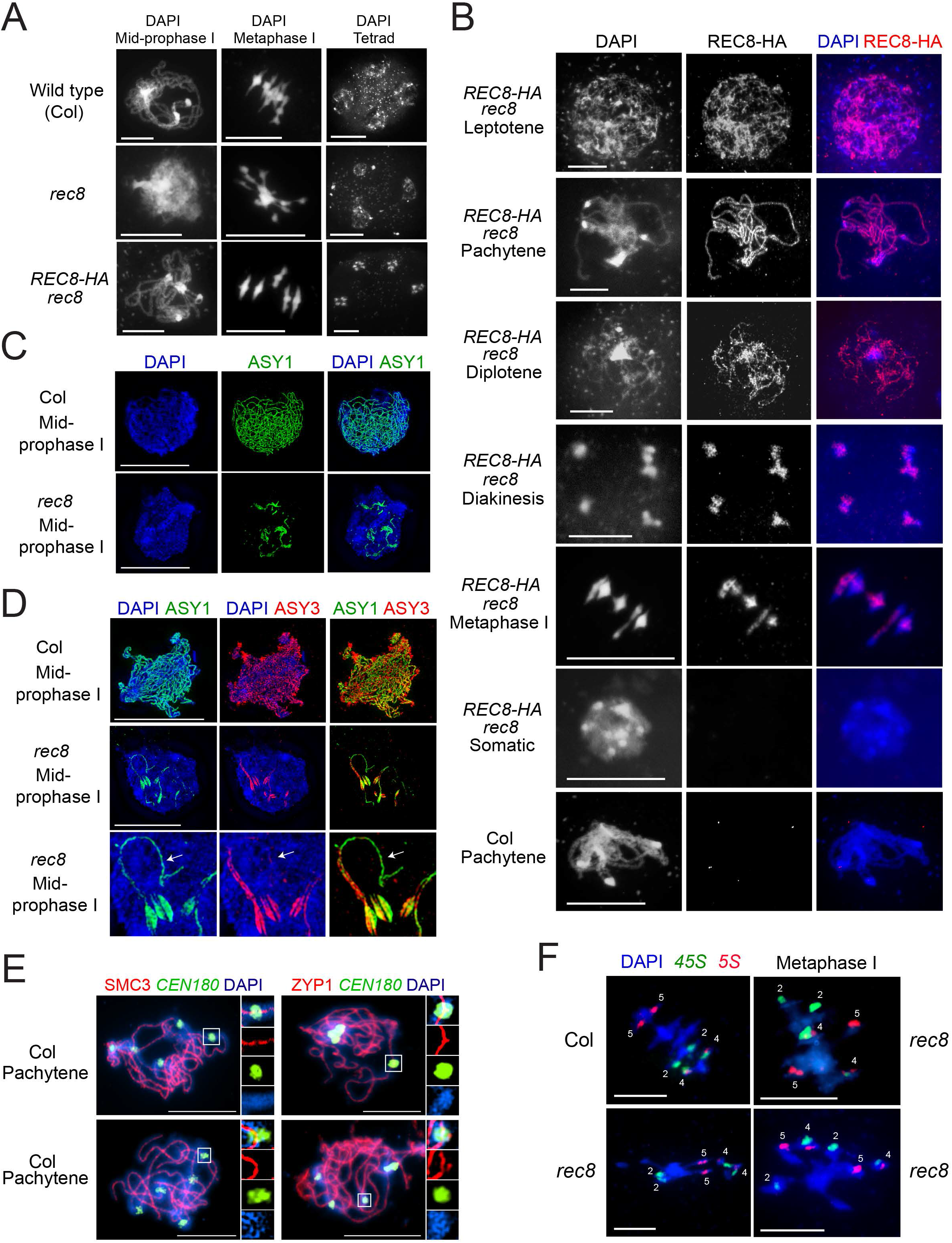
Complementation of *rec8* meiotic catastrophe via epitope tagging. (**A**) DAPI-stained spreads of wild type (Col), *rec8* and *REC8-HA rec8* male meiocytes at the labeled stages of meiosis. (**B**) Male meiocytes were stained for REC8-HA (red) and DAPI (blue), in the labeled genotypes and stages. (**C**) SIM images of wild type and *rec8* male meiocytes at early prophase I stained for ASY1 (green) and DAPI (blue). (**D**) As for C, but stained for ASY1 (green), ASY3 (red) and DAPI (blue). Close-ups of *rec8* axis polycomplexes are shown (lower). White arrows indicate regions staining for ASY1 but not ASY3. (**E**) Male meiocytes at pachytene stained for SMC3 (red) or ZYP1 (red), DAPI (blue) and *CEN180* FISH (green). Inset images show zooms of the *CEN180*-positive regions. (**F**) Male meiocytes at metaphase I with FISH performed against *45S* (green) and *5S* (red) rDNA. The positions of chromosomes 2, 4 and 5 are indicated. All scale bars=10 μm.

To analyze REC8 accumulation on chromosomes we immunostained male meiocytes and observed co-localization with chromatin from leptotene onwards, including within heterochromatin (Fig. 1B and Supplemental Fig. S2B-S2C). REC8 and chromatin co-localize as pairing and synapsis occurs, with strong co-staining at pachytene (Fig. 1B and Supplemental Fig. S2B-S2C). REC8 persists on bivalents through diakinesis and metaphase I (Fig. 1B and Supplemental Fig. S2B), as reported (Chelysheva et al. 2005; Cai et al. 2003). No signal was detected in non-transgenic wild type meiocytes, or somatic cells of the epitope-tagged lines (Fig. 1B and Supplemental Fig. S2B). We also performed western blotting on meiotic-stage flowers, which revealed bands of the expected size, in addition to bands with a ~20 kDa higher molecular mass (Supplemental Fig. S3). Similar heavy REC8 bands have been observed in yeast and mice and represent phosphorylated forms (Watanabe and Nurse 1999; Kitajima et al. 2003).

### REC8 is the major Arabidopsis kleisin required for meiotic axis polymerization and prevention of non-homologous recombination

Localization of cohesin and axis proteins are interdependent in many species (Kim et al. 2010; Severson et al. 2009; Storlazzi et al. 2008; Chelysheva et al. 2005). Therefore, we used immunocytology with epifluorescence and super-resolution structured illumination microscopy (SIM) to analyze Arabidopsis axis proteins ASY1 and ASY3 in *rec8* (Fig. 1C-1D). In wild type, ASY1 and ASY3 co-localize along linear axes from leptotene until zygotene (Fig. 1D) (Ferdous et al. 2012). In *rec8*, ASY1 and ASY3 occur in polycomplexes that persist through mid-prophase I (Fig. 1C-1D). The length of *rec8* ASY1 polycomplexes was shorter compared to the wild type axis at leptotene (mean=30 μm vs. 220 μm, MWW test *P*=3.02×10^-11^) (Supplemental Table S3). Heterochromatic chromocenters were observed within and apart from ASY1 polycomplexes in *rec8* (Supplemental Fig. S1C), suggesting they are independent structures. To analyze cohesin and the SC within the centromeres, we immunostained for SMC3 or ZYP1, in combination with fluorescence *in situ* hybridization (FISH) for the *CEN180* satellite repeats (Fig. 1E). At the zygotene–pachytene transition, we observed five *CEN180*-positive regions through which the SMC3 cohesin and ZYP1 SC signals were continuous and relatively uniform (Fig. 1E).

To investigate the *rec8* chromosome entanglements observed during prophase I, we performed FISH using *45S* and *5S* rDNA probes (Fig. 1F). This revealed physical connections between non-homologous chromosomes in *rec8*, not observed in wild type (Fig. 1F). Therefore, abnormal axis polymerization in *rec8* is associated with loss of recombination fidelity, leading to non-homologous joint molecules that resolve catastrophically at metaphase I (Fig. 1A and 1F). Although Arabidopsis encodes four α-kleisins (*REC8/SYN1, SYN2, SYN3* and *SYN4*), *REC8* shows strongest transcription in meiotic RNA-seq data (Supplemental Fig. S4A) (Walker et al. 2018). Furthermore, we measured crossovers in *syn2* and *syn4* mutants using chiasmata counting and fluorescent tagged lines (FTL), and did not observe significant differences to wild type (Supplemental Fig. S4B-S4D and Supplemental Tables S4-S6).

### REC8 is enriched in centromeric heterochromatin

To map REC8 localization throughout the genome we performed ChIP-seq using *REC8-HA rec8* or *REC8-Myc rec8* floral buds (Fig. 2A-2B and Supplemental S5), which contain all stages of meiosis. ChIP and input DNA were sequenced and log_2_(ChIP/input) enrichment calculated (Supplemental Table S7). Z-score standardization was applied so that the genome-wide mean equals zero and a value of one equates to one standard deviation from the mean. Biological replicate libraries for REC8-HA ChIP-seq were highly correlated (10 kb *r_s_*=0.99), as were REC8-HA and REC8-Myc (10 kb *r_s_*=0.81), at varying physical scales (Supplemental Fig. S5 and Supplemental Table S8), so REC8-HA data were used for subsequent analyses.

**Figure 2.**
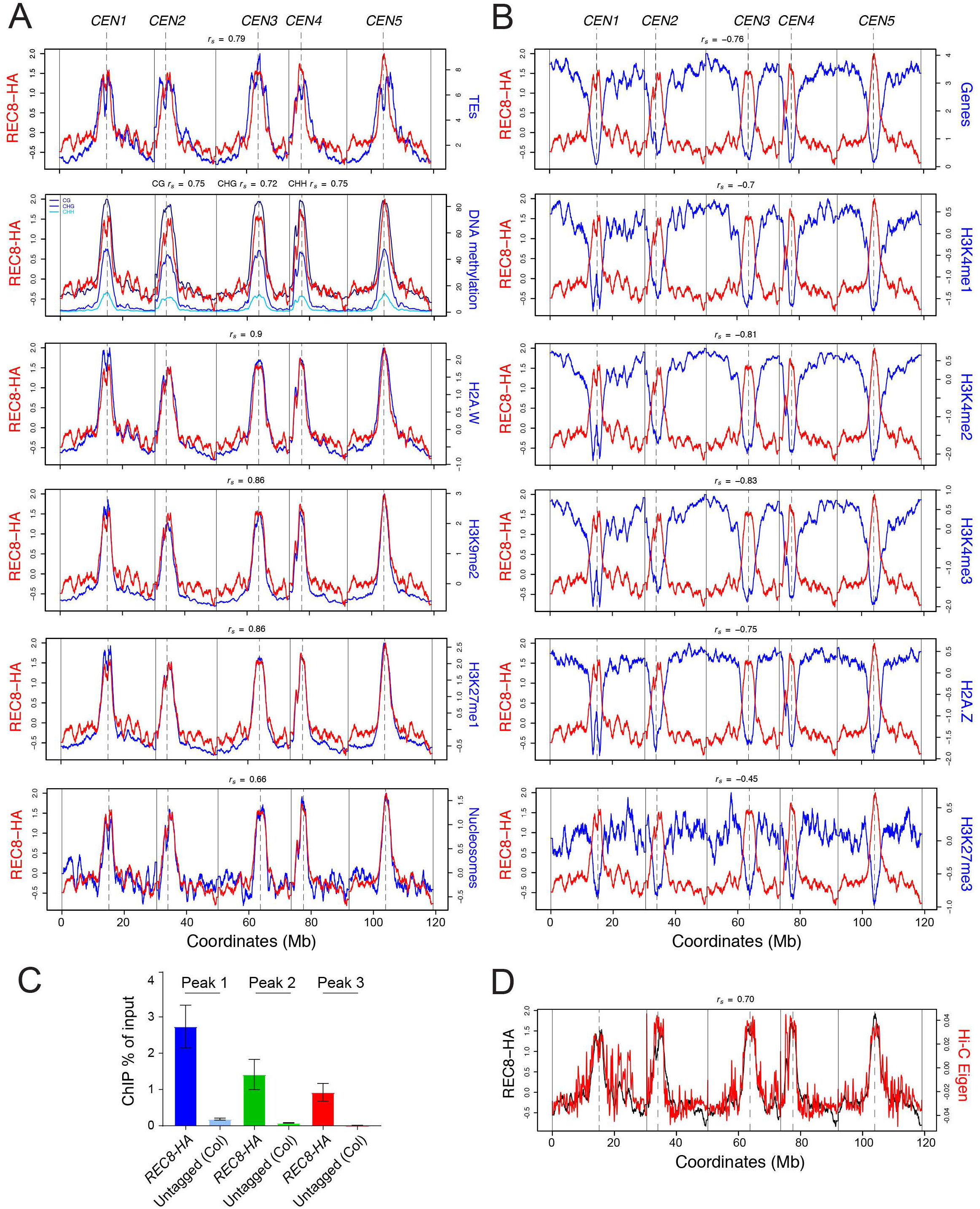
Genomic landscapes of REC8-cohesin, euchromatin and heterochromatin. (**A**) Genome-wide profiles (log_2_(ChIP/input) of REC8-HA (red) compared with transposable element density, DNA methylation (%) (Stroud et al. 2013), H2A.W (Yelagandula et al. 2014), H3K9me2, H3K27me1 and nucleosomes (blue) (Choi et al. 2018). Vertical solid lines indicate telomeres and dotted lines indicate centromeres. Spearman’s correlation coefficients (*r_s_*) are printed above the plots. (**B**) As for A, but plotting gene density, H3K4me1, H3K4me2, H3K4me3 (Choi et al. 2018), H2A.Z (Yelagandula et al. 2014) and H3K27me3 (blue). (**C**) ChIP-qPCR enrichment (% input) in *REC8-HA rec8* and untagged wild type (Col) floral buds at REC8 peaks. (**D**) REC8-HA ChIP-seq data (black) plotted against Hi-C Eigenvalues (red) that correspond to the first principle component of the contact matrix, where the sign of the value denotes compartment (Liu et al. 2016).

At the chromosome scale, REC8 enrichment was greatest in the centromeres and in proximal heterochromatin (Fig. 2A-2B). Consequently, we observed positive correlations between REC8 and transposable elements (*r_s_*=0.79), DNA methylation in CG (*r_s_*=0.75), CHG (*r_s_*=0.72) and CHH (*r_s_*=0.75) sequence contexts and the heterochromatic histone modifications H3K9me2 (*r_s_*=0.86), H3K27me1 (*r_s_*=0.86) and histone variant H2A.W (*r_s_*=0.90) (Fig. 2A, Supplemental Fig. S6 and Supplemental Table S9) (Stroud et al. 2013; Yelagandula et al. 2014). Nucleosomes, measured via MNase-seq, also show high pericentromeric enrichment and a positive correlation with REC8 (*r_s_*=0.66) (Fig. 2A). In contrast, genome-wide negative correlations between REC8 and gene density (*r_s_*=-0.76), and the gene-associated chromatin modifications H3K4me1 (*r_s_*=-0.70), H3K4me2 (*r_s_*=-0.81), H3K4me3 (*r_s_*=-0.83), H2A.Z (*r_s_*=-0.75) and H3K27me3 (*r_s_*=-0.45) were observed (Fig. 2B, Supplemental Fig. S6 and Supplemental Table S9) (Yelagandula et al. 2014). Hence, at the chromosome scale, REC8 enrichment is strongly correlated with heterochromatin, although substantial signal occurs in the gene-rich chromosome arms. To validate ChIP specificity we analyzed REC8 peak loci using qPCR in *REC8-HA rec8* compared to untagged wild type, and observed significant enrichment in all cases (Fig. 2C and Supplemental Table S10).

Hi-C studies have revealed A/B compartment organization in plant genomes, where the A compartment contains gene-rich euchromatin and the B compartment contains the centromeres and heterochromatin (Feng et al. 2014; Liu et al. 2016). We compared REC8 ChIP-seq data to a high resolution Hi-C map generated from Arabidopsis seedlings (Fig. 2D) (Liu et al. 2016). Hi-C Eigenvalues correspond to the first principle component of the contact matrix, where the sign of the value denotes the compartment (negative=A, positive=B) (Liu et al. 2016). Hi-C Eigenvalues showed a strong positive correlation with REC8 (*r_s_*=0.70) (Fig. 2D), indicating enrichment in the B compartment. We compared REC8 ChIP enrichment with cytogenetic maps made at pachytene that relate meiotic axis length (μm) to physical DNA length (kb) (Fransz et al. 2000). For example, the chromosome 4 heterochromatic knob contains 614 kb/μm, compared to euchromatic regions with a mean of 356 kb/μm (Supplemental Table S11) (Fransz et al. 2000). Differential condensation of these regions is reflected by a 43-fold difference in REC8 ChIP-seq enrichment (Supplemental Table S11). Therefore, A/B compartment structure is likely a dominant feature of both mitotic and meiotic chromosomes in Arabidopsis, consistent with pronounced heterochromatin differentiation at pachytene (Fig. 1A-1B) (Fransz et al. 2000).

### REC8 anti-correlates with meiotic DSBs and crossovers

We investigated the relationships between REC8, meiotic DSBs mapped using SPO11-1-oligos, and 3,320 crossovers mapped by genotyping-by-sequencing of Col×Ler F_2_ individuals (Choi et al. 2018; Underwood et al. 2018). At the chromosome scale, SPO11-1-oligos and crossovers show negative correlations with REC8 (*r_s_*=-0.22 and *r_s_*=-0.15), and these relationships are strengthened in the pericentromeres (*r_s_*=-0.94 and *r_s_*=-0.84) (Fig. 3A and Supplemental Fig. S6). However, variation in the ratio of SPO11-1-oligos, crossovers and REC8 was observed along the chromosomes (Fig. 3A). Multiple factors likely contribute to this variation, including patterns of interhomolog polymorphism, chromatin states and the action of crossover interference.

**Figure 3.**
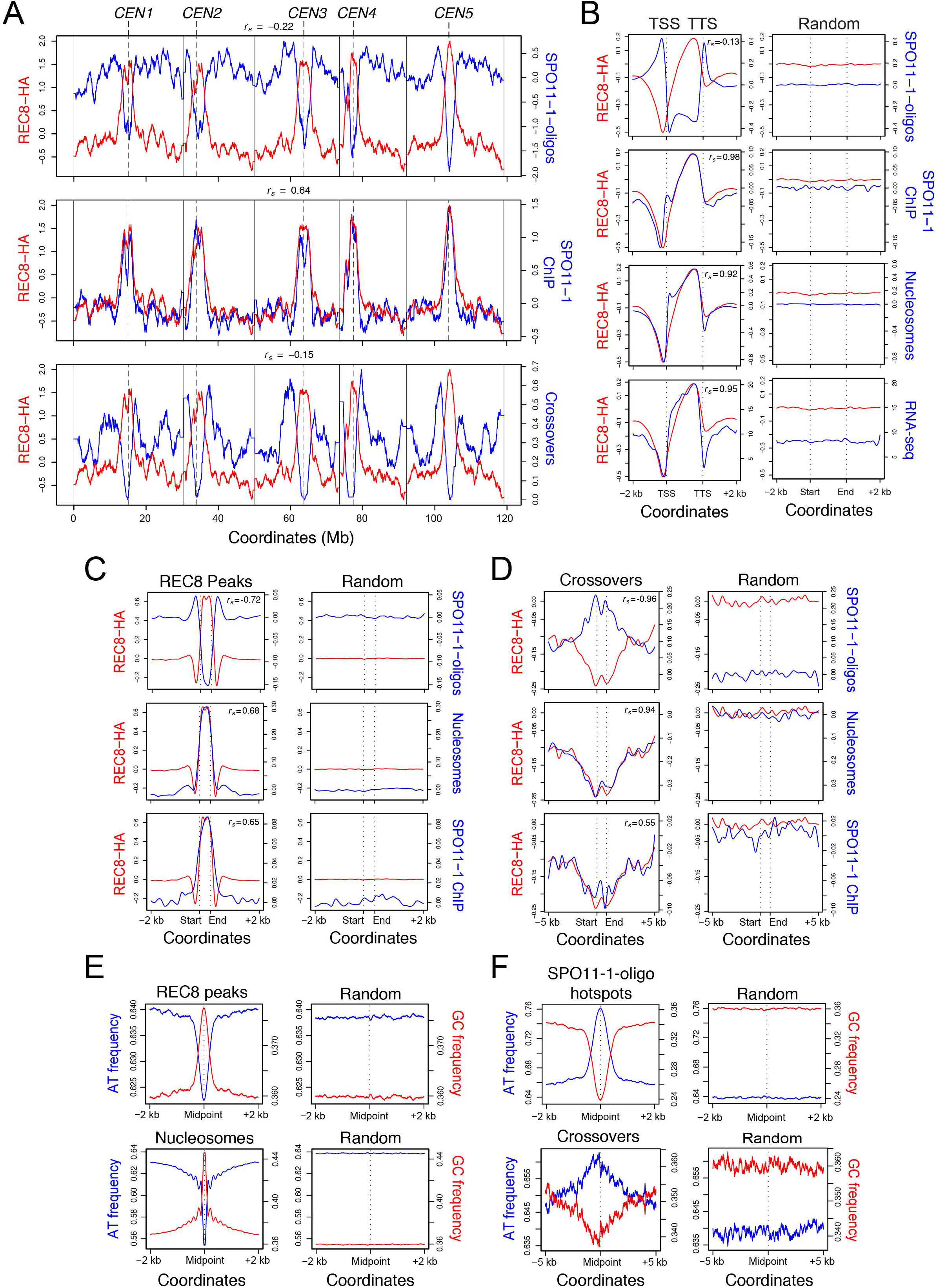
REC8 enrichment correlates with suppression of meiotic DSBs and crossovers. (**A**) Genome-wide profiles (log_2_(ChIP/input) of REC8-HA (red) compared with SPO11-1-oligos (Choi et al. 2018), SPO11-1 ChIP and crossovers (blue) (Choi et al. 2018). Vertical solid lines indicate telomeres and dotted lines indicate centromeres. Spearman’s correlation coefficients (*r_s_*) are printed above. (**B**) REC8-HA (red), SPO11-1-oligos, SPO11-1 ChIP-seq, RNA-seq and nucleosomes (blue) plotted between gene transcriptional start and termination sites (TSS-TTS) and 2 kb flanking regions, compared to the same number of random regions of the same widths. (**C**) Average coverage profiles of REC8-HA (red) compared with SPO11-1-oligos, SPO11-1 ChIP and nucleosomes (blue) within REC8-HA peaks, compared to the same number of random positions of the same widths. (**D**) As for C, but plotting over crossover intervals and 5 kb flanking regions, compared to the same number of random positions. Mean resolved crossover widths are shown by dotted lines. (**E**) Base frequencies (A+T=blue; G+C=red) plotted in 2 kb regions around the midpoints of REC8 peaks or nucleosomes and compared to random positions. (**F**) As for E, but analyzing SPO11-1-oligo hotspots with 2 kb of flanking sequence, or crossovers with 5 kb.

We previously observed SPO11-1 immunostaining in association with ASY1 throughout prophase I (Choi et al. 2018). To provide further insight into SPO11-1 binding to the genome we performed ChIP-seq, using the SPO11-1-Myc line previously used for SPO11-1-oligo sequencing (Supplemental Table S9) (Choi et al. 2018). At the chromosome scale, SPO11-1 ChIP signal showed a positive correlation with REC8 (*r_s_*=0.64) and nucleosomes (*r_s_*=0.87), and a negative correlation with SPO11-1-oligos (*r_s_*=-0.69) and crossovers (*r_s_*=-0.50) (Fig. 3A). Hence, while SPO11-1-oligos capture the DSB landscape, crosslinking of SPO11-1 to chromatin throughout meiosis reveals an axis-associated signal similar to REC8. These data are consistent with recruitment of chromatin loops to axis-associated SPO11-1 during DSB formation and repair (Blat et al. 2002; Panizza et al. 2011).

At the fine scale, Arabidopsis shows SPO11-1-oligo enrichment in nucleosome-depleted gene promoters and terminators (Fig. 3B) (Choi et al. 2018). In contrast, REC8 and SPO11-1 ChIP-seq show strongest enrichment within gene bodies towards transcriptional termination sites (TTSs), and were positively correlated with one another (*r_s_*=0.98) (Fig. 3B). The REC8 profile within genes also strikingly correlates with transcript abundance and nucleosomes (*r_s_*=0.95 and *r_s_*=0.92) (Fig. 3B). We analyzed introns and exons, orientated in gene 5′ to 3′ directions, and observed that SPO11-1-oligos were relatively depleted from exons and enriched within introns, whereas REC8, SPO11-1 ChIP, transcripts and nucleosomes show the opposite trends (Supplemental Fig. S7). This reveals fine-scale variation in the meiotic axis and recombination machinery in relation to gene organization, transcription and chromatin.

To further analyze REC8 at the fine scale, we identified 87,738 peaks with a mean width of 475 bp (Fig. 3C, Supplemental Fig. S8A and Supplemental Table S12). Consistent with the trends at the chromosome scale and within genes, REC8 peak enrichment positively correlated with nucleosomes (*r_s_*=0.68) and SPO11-1 ChIP (*r_s_*=0.65), but negatively with SPO11-1-oligos (*r_s_*=-0.72) (Fig. 3C). We applied permutation tests and observed that significantly fewer-than-expected REC8 peaks overlap SPO11-1-oligo hotspots and crossovers, and more-than-expected REC8 peaks overlap nucleosome positions and SPO11-1 ChIP peaks (all P<0.0001) (Supplemental Fig. S9 and Supplemental Table S12). We analyzed crossover intervals and 5 kb flanking regions and observed that REC8, nucleosomes and SPO11-1 ChIP were all depleted, whereas SPO11-1-oligos were enriched (Fig. 3D and Supplemental Fig. S8B). Finally, we investigated DNA sequence composition and observed that REC8 peaks and well-positioned nucleosomes show a strong GC bias, whereas SPO11-1-oligo hotspots and crossovers are AT-biased (Fig. 3E-3F). Therefore, REC8 enrichment at both the chromosome and the fine scale is associated with suppression of SPO11-1-oligos and crossovers, and correlates with the presence of heterochromatin and GC-rich DNA sequences.

### REC8-cohesin and meiotic DSB landscapes are remodeled in *kyp suvh5 suvh6* H3K9me2 mutants

In plants, non-CG DNA methylation and H3K9me2 maintain each other in a self-reinforcing epigenetic loop (Stroud et al. 2014, 2013). For example, the *kryptonite/suvh4 suvh5 suvh6 (kss)* SET domain triple mutant eliminates both H3K9me2 and non-CG DNA methylation (Stroud et al. 2014, 2013). Mutants in the non-CG/H3K9me2 pathway also show increased pericentromeric DSBs and crossovers in Arabidopsis (Underwood et al. 2018). Therefore, we performed REC8-HA ChIP-seq in *kss* mutants to test for interactions between cohesin, chromatin and recombination. At the chromosome scale, the *kss* mutant shows slight but significant changes in REC8 ChIP enrichment, with decreases in the centromeric heterochromatin (one-tailed MWW test *P*=0.008), and increases in the chromosome arms (one-tailed MWW test *P*=0.001) (Supplemental Fig. S10). We performed REC8-HA immunostaining in wild type and *kss* and observed a slight decrease in signal over the heterochromatin, although this was not significant (Supplemental Tables S13-S14). Furthermore, the level of REC8 loading in *kss* is sufficient to maintain sister chromosome cohesion and no decrease in pollen viability occurs, compared to wild type (MWW test *P*=0.29) (Fig. 4A and Supplemental Table S15). Hence, REC8 function in maintaining cohesion during meiosis is not H3K9me2-dependent in Arabidopsis. To assess chromatin compaction during meiosis, pachytene cells were DAPI-stained and the proportion of area occupied by heterochromatin measured. A slight but significant decrease in heterochromatic compaction was observed in *kss* (MWW test *P*=0.047) (Supplemental Table S16), which is consistent with changes to Hi-C contact maps in *kss* heterochromatin (Feng et al. 2014).

**Figure 4.**
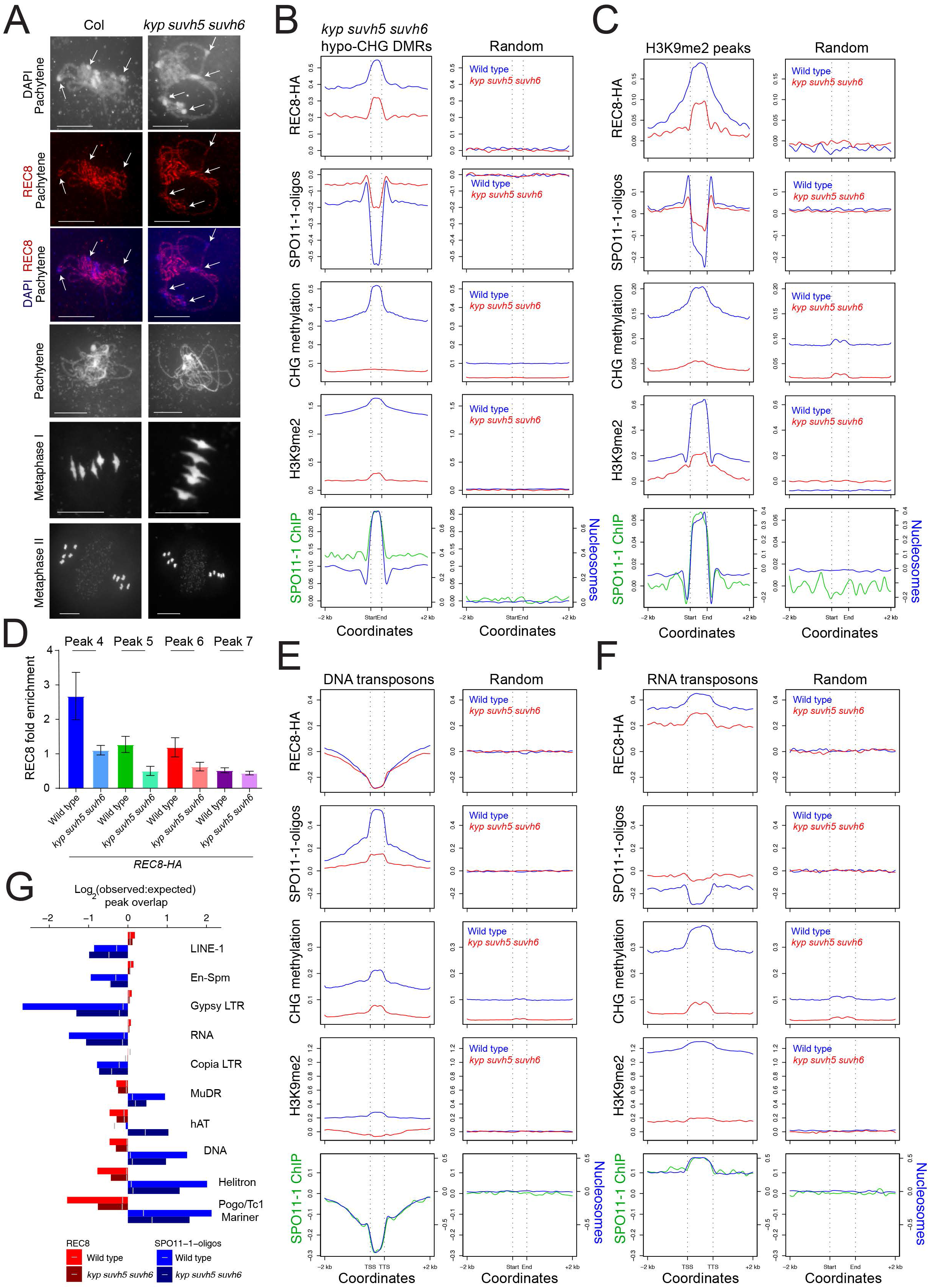
REC8, chromatin and meiotic DSB landscapes are remodeled in *kyp suvh5 suvh6*. (**A**) DAPI-stained or HA immunostained (red) spreads of wild type and *kss* male meiocytes at the labeled stages. White arrows indicate chromocenters. All scale bars=10μm. (**B**) REC8-HA (log_2_(ChIP/input)), SPO11-1-oligos, CHG methylation and H3K9me2 from wild type (blue) and *kss* (red) within hypo-CHG DMRs (Stroud et al. 2013), and 2 kb flanking regions, or the same number of random positions of the same widths. The lower plot shows nucleosomes and SPO11-1 ChIP-seq in wild type within the hypo-CHG DMRs. (**C**) As for B, but plotting in relation to H3K9me2 peaks identified in wild type. (**D**) Fold change of REC8 ChIP-qPCR enrichment at REC8 peaks in *REC8-HA rec8* compared to *kss*, normalized by Peak 3 which has low H3K9me2 and did not change in *kss*. Peak 7 is a negative control region not expected to change in *kss*. (**E**) As for B, but plotting in relation to DNA transposons. (**F**) As for B, but plotting in relation to RNA transposons. (**G**) Bar graphs showing permutation test derived log_2_(observed:expected) overlap of REC8 and SPO11-1-oligo peaks in wild type and *kss* with different transposon families. Vertical gray lines mark significance thresholds (a=0.05).

We next analyzed REC8, chromatin and recombination in wild type versus *kss* mutants at the fine scale. For example, 15,562 hypo-CHG differentially DNA methylated regions (DMRs) were previously identified in *kss* (Stroud et al. 2013), which in wild type are nucleosome-enriched and DSB-suppressed (Fig. 4B). We observed that these hypo-CHG DMRs lose both REC8 and H3K9me2, and gain SPO11-1-oligos in *kss* (one-tailed MWW tests, all *P*<2.2×10^-16^) (Fig. 4B). As a further test we analyzed H3K9me2 peaks defined in wild type (n=20,289), which showed a similar pattern to hypo-CHG DMRs, with coincident loss of H3K9me2, non-CG DNA methylation and REC8 and gain of SPO11-1-oligos in *kss* (all *P*<2.2×10^-16^) (Fig. 4C). We used ChIP-qPCR to analyze enrichment at REC8 peaks observed to lose both REC8 and H3K9me2 in *kss* (Fig. 4D and Supplemental Table S17). Using independent ChIP replicates this assay confirmed reduction of REC8 at the peak loci in *kss*, compared to a control locus that did not change (Fig. 4D and Supplemental Table S17).

We previously observed that DNA and RNA transposons are differentiated by chromatin state and levels of meiotic recombination in Arabidopsis (Choi et al. 2018). Compared to DNA elements, RNA elements show higher REC8 and nucleosomes, and lower SPO11-1-oligos (Fig. 4E-4G and Supplemental Fig. S11) (Choi et al. 2018). RNA elements also show higher H3K9me2 and non-CG methylation levels in wild type, which are reduced in *kss* (Fig. 4E-4F). As observed at hypo-CHG DMRs, loss of heterochromatic marks at RNA elements correlates with reduced REC8 and gain of SPO11-1-oligos in *kss* (Fig. 4E-4F). In contrast, DNA elements also show loss of heterochromatic marks in *kss*, while REC8 was unchanged (Fig. 4E-4G). Hence, REC8, recombination and chromatin are differentiated across transposon classes, with greatest cohesin accumulation on heterochromatic RNA elements that are highly enriched in proximity to the centromeres.

### REC8 enrichment in genes anti-correlates with transcription levels

Cohesin occupancy on chromosomes is strongly influenced by transcription (Busslinger et al. 2017; Lengronne et al. 2004; Misulovin et al. 2008; Kagey et al. 2010; Sun et al. 2015). Therefore, we compared REC8 enrichment with RNA-seq data generated from wild type meiotic-stage floral buds, or directly from male meiocytes (Fig. 5A) (Walker et al. 2018; Choi et al. 2018). In both cases we observed an anticorrelation between RNA expression and REC8 enrichment at the chromosome scale (*r_s_*=-0.80 and *r_s_*=-0.66) (Fig. 5A). Within gene bodies, REC8 enrichment was spatially correlated with RNA expression (*r_s_*=0.95), H3K4me1 (*r_s_*=0.87) and CG DNA methylation (*r_s_*=0.95), whereas negative correlations were observed between REC8 and chromatin modifications enriched at gene 5' ends, including H3K4me3 (*r_s_*=-0.64), H3K4me2 (*r_s_*=-0.49) and H2A.Z (*r_s_*=-0.64) (Fig. 5B). Therefore, REC8 accumulates within gene bodies, spatially coincident with chromatin marks associated with active transcription (H3K4me1 and CG methylation).

**Figure 5.**
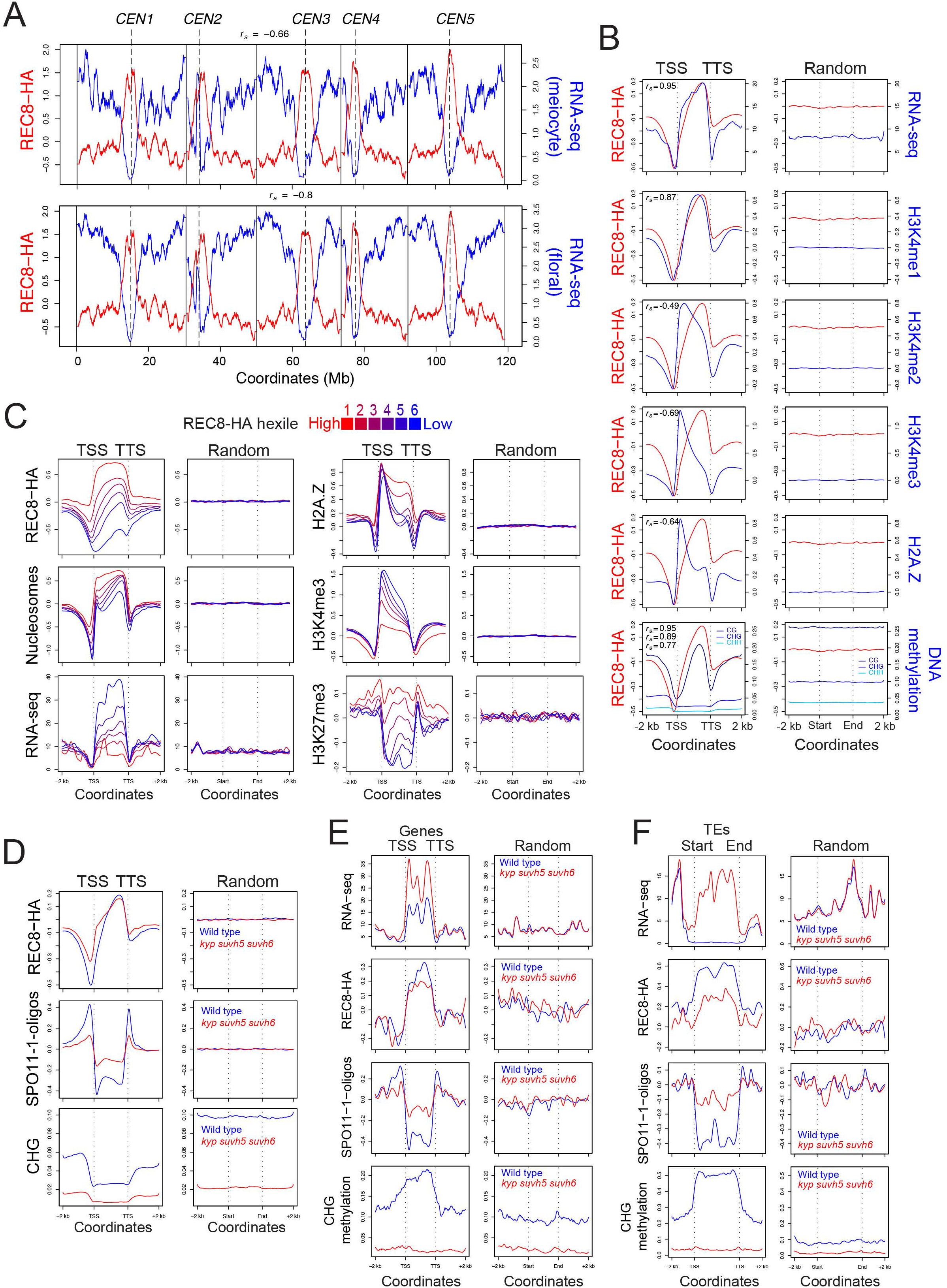
REC8 is shaped by chromatin state and transcription within genes and transposons. (**A**) REC8-HA (log_2_(ChIP/input) red) compared with RNA-seq from meiocytes and floral buds (blue) (Walker et al. 2018; Choi et al. 2018). Vertical solid lines indicate telomeres and dotted lines indicate centromeres. Spearman’s correlation coefficients (*r_s_*) are printed above the plots. (**B**) REC8-HA (red) was compared with RNA-seq (Choi et al. 2018), H3K4me1, H3K4me2, H3K4me3, H2A.Z and DNA methylation in CG, CHG and CHH sequence contexts (blue) (Stroud et al. 2013) in gene TSS-TTS and 2 kb flanking regions, or the same number of random regions of the same widths. Spearman’s correlation coefficients (*rs*) are printed above. (**C**) As for B, but analyzing the indicated parameters within genes that were ranked into hexiles according to REC8 levels between TSS and TTS (red=highest, blue=lowest). (**D**) As for B, but analyzing REC8-HA, SPO11-1-oligos or CHG DNA methylation between gene TSS and TTS, or the same number of random positions of the same widths, in wild type (blue) versus *kss* (red). (**E**) As for D, but analyzing genes that are transcriptionally upregulated in *kss* and plotting REC8-HA, SPO11-1-oligos and CHG DNA methylation. (**F**) As for E, but analyzing transposable elements (TEs) that are transcriptionally upregulated in *kss*.

To further investigate the relationship between transcription levels and REC8, we ranked genes into six groups (hexiles), according to REC8 enrichment within transcribed regions (TSSs-TTSs) (Fig. 5C and Supplemental Fig. S12). Average REC8 and transcription levels were negatively correlated (*r_s_*=-0.44), with highly transcribed genes showing lowest REC8 (Fig. 5C). Reciprocal patterns were observed when genes were ranked according to transcription, with low-expression genes showing highest REC8 (Supplemental Fig. S13). In addition, polarized REC8 enrichment occurs towards the TTS of gene hexiles with highest transcription (Fig. 5C), consistent with RNA polymerase pushing or evicting cohesin along transcribed genes. Gene hexiles with highest REC8 also show highest nucleosomes and H3K27me3 levels, lowest H3K4me3 and H2A.Z spreading throughout the gene body (Fig. 5C), which are features of facultatively silent genes in plants (Mozgova and Hennig 2015). Interestingly, analysis of H3K27me3 in relation to REC8 peaks showed enrichment in flanking positions, although the peaks themselves were depleted of this mark (Supplemental Fig. S8A). In conclusion, we observe intragenic REC8 enrichment, with highest levels in genes that are transcriptionally silenced by the Polycomb system and H3K27me3.

We performed RNA-seq from wild type and *kss* floral buds and compared these data to REC8, SPO11-1-oligo and DNA methylation data (Fig. 5D). Genes in wild type show low levels of CHG DNA methylation, which was further reduced in *kss* (Fig. 5D). We observed that gene promoters and terminators show increased REC8 and reduced SPO11-1-oligos in *kss*, whereas opposite trends are observed within gene bodies (Fig. 5E). We performed differential expression analysis and identified 179 transposable elements (TEs) and 263 genes that were transcriptionally upregulated in *kss* (FDR<0.01) (Supplemental Fig. S14 and Supplemental Table S18). We observed reduced REC8 and gain of SPO11-1-oligos over the upregulated genes and TEs, coincident with loss of CHG DNA methylation (Fig. 5E-5F). These loci provide examples of local changes to chromatin state and transcription that influence REC8 occupancy and recombination, within both genes and transposons.

### Axis polycomplexes in *rec8* recruit the homologous recombination machinery with altered stoichiometry

Our genomic data revealed that REC8 accumulates in regions of low meiotic DSBs and crossovers in Arabidopsis. We were therefore interested to further investigate the cause of meiotic chromosome fragmentation in *rec8*. As we observed evidence for non-homologous recombination in *rec8* using FISH (Fig. 1F), we hypothesized that *rec8* axis polycomplexes may recruit the recombination machinery. To visualize meiotic DSBs, we immunostained male meiocytes for γH2A.X and observed a mean of 202 axis foci in wild type (Fig. 6A and Supplemental Table S19). In *rec8*, γH2A.X foci were significantly reduced (mean=53, MWW test *P*=3.37×10^-6^), although they remained associated with ASY1 polycomplexes (Fig. 6A and Table S19). We immunostained for the ssDNA binding proteins RPA1a, RAD51 and DMC1, which show mean axis-associated foci numbers of 181, 174 and 172 respectively at mid-prophase I in wild type, which were significantly reduced in *rec8* to 39, 36 and 14 (MWW tests, *P*=3.37×10^-6^, *P*=3.33×10^-6^, *P*=1.08×10^-5^) (Fig. 6B-6C, Supplemental Fig. S15A and Supplemental Tables S19-S20). We also immunostained for SPO11-1-Myc and observed foci distributed throughout the nucleus that were axis-associated at leptotene and persisted until pachytene (Supplemental Fig. S15B and Supplemental Table S21). In *rec8*, SPO11-1-Myc signal was distributed throughout the nucleus, both within and apart from ASY1 polycomplexes (Supplemental Fig. S15B). The presence of SPO11-1 in regions lacking axis polycomplexes in *rec8*, which also show an absence of DSB markers, is consistent with the axis being required to promote DSBs (Supplemental Fig. S15B). In support of this, a positive correlation exists between axis length and γH2A.X and RAD51 foci between nuclei, in both wild type and *rec8* (wild type *r_s_*=0.83 and 0.69, *rec8 r_s_*=0.92 and 0.77) (Supplemental Table S22). Together, this provides cytological evidence that DSB formation and interhomolog strand invasion are associated with *rec8* axis polycomplexes.

**Figure 6.**
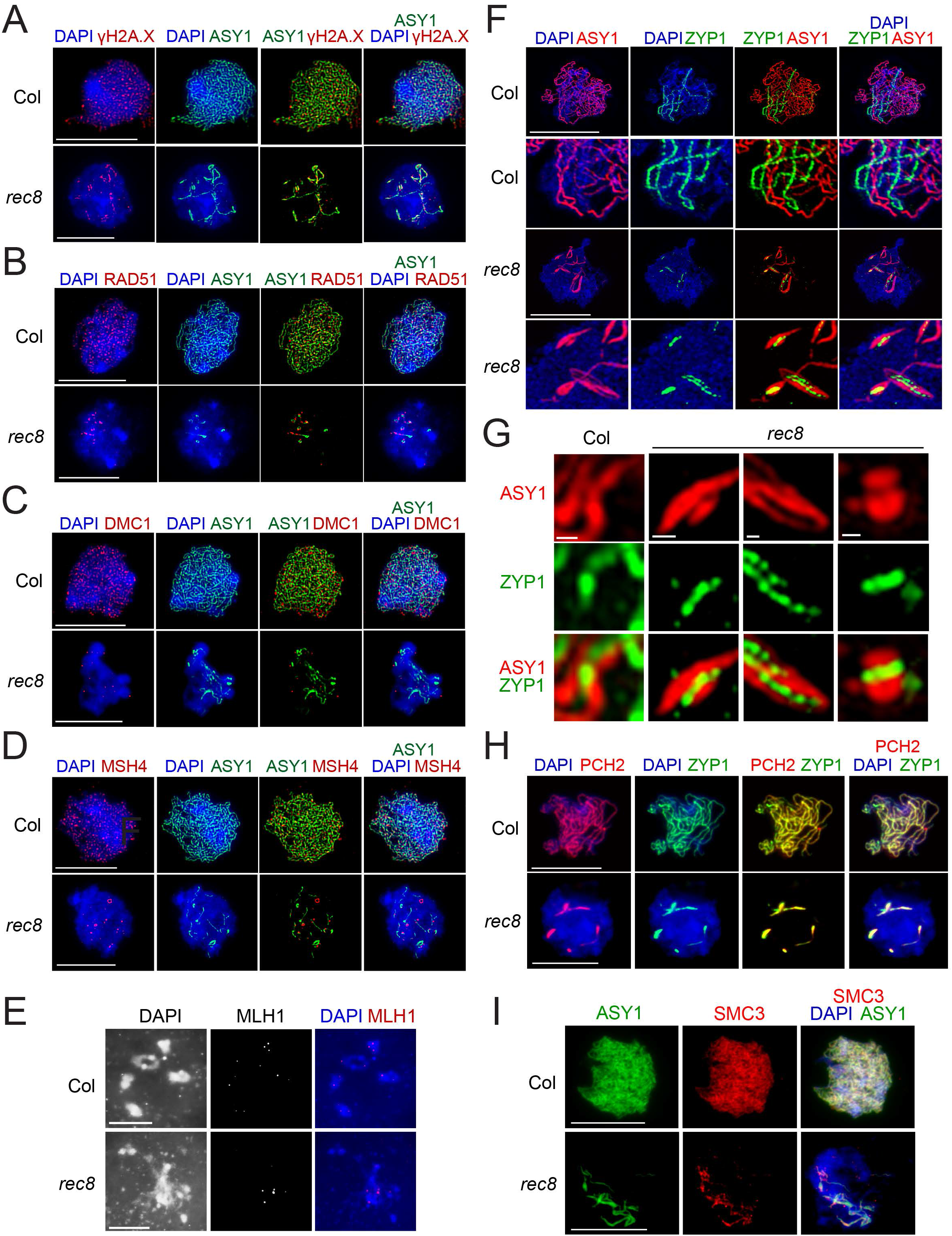
Axis polycomplexes recruit the homologous recombination machinery with altered stoichiometry in *rec8*. (**A**) Wild type and *rec8* male meiocytes in midprophase I were stained for ASY1 (green), yH2A.X (red) and DAPI (blue). In parts A-D, wild type nuclei are at leptotene. (**B**) As for A, but staining for RAD51 (red) and ASY1 (green). (**C**) As for A, but staining for DMC1 (red) and ASY1 (green). (**D**) As for A, but staining for MSH4 (red) and ASY1 (green). (**E**) As for A, but at diakinesis stage and stained for MLH1 (red) and DAPI (blue). (**F**) SIM images of male meiocytes at zygotene stage stained for ASY1 (red), ZYP1 (green) and DAPI (blue). Close-ups of synapsed regions are shown in the lower panels, for each genotype. (**G**) As for F, but showing close-ups of ASY1 (red) and ZYP1 (green) staining. Scale bars=200 nm. (**H**) As for F, but staining for PCH2 (red), ZYP1 (green) and DAPI (blue). (**I**) Male meiocytes in midprophase I stained for ASY1 (green), SMC3 (red) and dApI (blue). All scale bars=10 μm, apart from in (G).

We next immunostained for ZMM repair factors, which are required for formation of interfering crossovers (Hunter 2015). In wild type, the MutS homolog MSH4 forms a mean of 179 axis-associated foci at leptotene, which were significantly reduced in *rec8* (mean=13 foci, MWW test *P*=1.10×10^-5^) (Fig. 6D and Supplemental Table S20). The MutL homolog MLH1 acts late in prophase I and shows a mean of 10.4 chiasmata-associated foci at diakinesis, which were also significantly reduced in *rec8*, yet remained associated with the entangled chromosomes at late prophase I (mean=4.9 foci, MWW test *P*=2.59×10^-10^) (Fig. 6E and Supplemental Table S23). It is notable that MSH4 and DMC1 foci showed a greater reduction in foci numbers (7.3% and 8.3% of wild type), compared with γH2A.X, RAD51, RPA1a and MLH1 (26.2%, 19.9%, 22.4% and 47% of wild type) (Supplemental Fig. S15C and Supplemental Tables S19-S20 and S23). Therefore, although *rec8* polycomplexes recruit recombination foci, they are reduced in number and show altered stoichiometry relative to wild type. We propose that a subset of these foci represent non-homologous recombination events in *rec8* that lead to joint molecules and fragmentation at metaphase I.

We used immunocytology with epifluorescence and SIM to analyze Arabidopsis axis (ASY1, ASY3 and SMC3) and SC (ZYP1) components in wild type and *rec8* (Fig. 6F-6I). Chromosome synapsis initiates at zygotene with the formation of ZYP1 stretches, which become depleted of ASY1 (Fig. 6F), until full synapsis is achieved at pachytene (Ferdous et al. 2012). We observed short stretches of ZYP1 polymerization between ASY1 polycomplexes in *rec8* (Fig. 6F). In wild type ZYP1 polymerizes between axes separated by a mean distance of 109 nm (Fig. 6G and Supplemental Table S24), consistent with eukaryotic SC widths (Zickler and Kleckner 1999). In *rec8*, ZYP1 was detected between ASY1 polycomplexes, with a mean distance not significantly different from wild type (119 nm, MWW test *P*=0.22) (Fig. 6G and Supplemental Table S24). PCH2 is a conserved meiotic AAA+ATPase required to remodel the axis during synapsis, which forms a linear signal with ZYP1 at pachytene (Fig. 6H) (Lambing et al. 2015). The *rec8* polycomplexes co-stained for both PCH2 and ZYP1 (Fig. 6H). Interestingly, the SMC3 cohesin subunit was recruited to ASY1 polycomplexes, despite the absence of REC8 (Fig. 6I). Therefore, *rec8* polycomplexes include ASY1, ASY3 and SMC3 and can recruit PCH2 and ZYP1 to produce synapsed structures with a similar inter-axis width to wild type. These cytological data support a role for REC8-cohesin in organizing correct polymerization of the axis and SC and promoting high-fidelity interhomolog recombination.

## Discussion

During meiosis, replicated sister chromatids are organized as linear loop arrays connected to the axis, where REC8 is enriched (Zickler and Kleckner 1999; Blat et al. 2002; Panizza et al. 2011). Comparative analysis across eukaryotes supports a conserved density of ~20 chromatin loops per μm of axis at pachytene, with larger genomes having increased axis length and/or chromatin loop size (Zickler and Kleckner 1999). The 125 Mb Arabidopsis genome has an axis length of 331 μm equating to 378 kb/μm (Fransz et al. 2000). Assuming 20 loops per μm, this gives an estimate of 18.9 kb per loop (Fig. 7). Cytogenetic maps have also revealed that heterochromatin is more condensed (614 kb/μm) than euchromatic regions (356 kb/μm) at pachytene (Fransz et al. 2000), equating to loop size estimates of 30.7 kb and 17.8 kb, respectively (Fig. 7). However, Arabidopsis meiotic chromosome spreads show that chromatin loops extend ~300 nm from the axis, which does not vary between euchromatin and heterochromatin (Ferdous et al. 2012; Armstrong et al. 2002). Therefore, to accommodate additional sequence per μm of axis, the heterochromatic loops must be more highly condensed (Fig. 7). This may enhance REC8 crosslinking to heterochromatin during ChIP, and is also reflected by interphase chromocenter organization and B-compartment identity (Liu et al. 2016; Feng et al. 2014; Fransz et al. 2000). It is also important to note that REC8 is lost from the chromosome arms at ~30 hours post S-phase, whereas it persists in the centromeres until ~33 hours (Cai et al. 2003; Chelysheva et al. 2005). As we sample over all meiotic stages, this may further contribute to relative centromeric REC8 ChIP enrichment.

**Figure 7.**
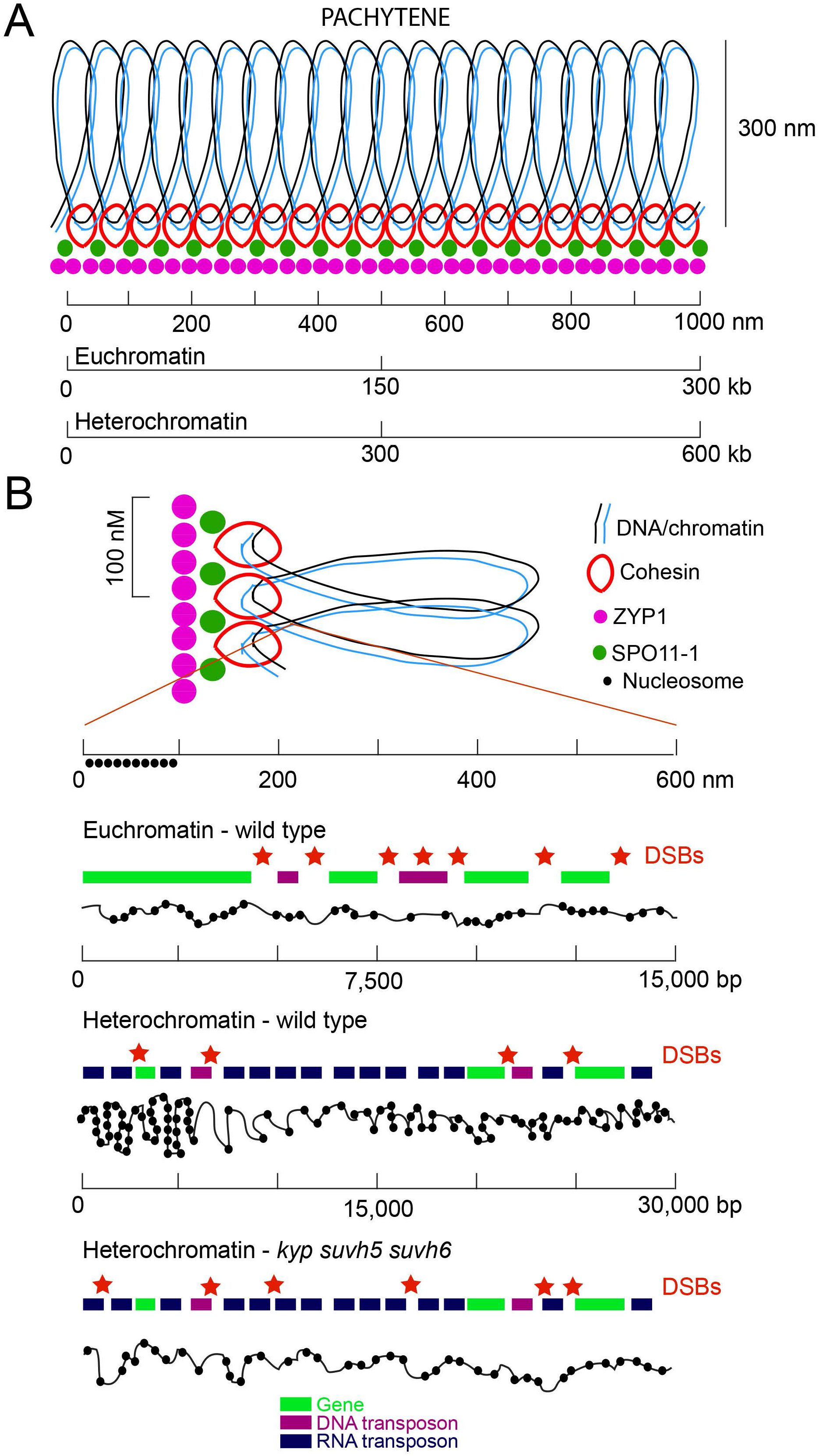
Cohesin and chromatin orchestrate meiotic chromosome architecture and interhomolog recombination. (**A**) Linear loop arrays of sister chromatids (black and blue) are tethered to the meiotic axis at pachytene stage, via passage through REC8-cohesin rings (red). Associated with cohesin are SPO11-1 (green) and the synaptonemal complex protein ZYP1 (purple). Chromatin loops extend from the axis for ~300 nm (Ferdous et al. 2012; Armstrong et al. 2002). A 1 μm section of axis is shown with 20 loops (Zickler and Kleckner 1999), with the physical length of DNA associated with 1 μm of axis inferred from FISH experiments (Fransz et al. 2000). (**B**) A close-up of A is shown, with one loop beneath corresponding to 600 nm and ~15 kb in euchromatin and ~30 kb in heterochromatin. Chromatin fiber compaction is shown by the black line with nucleosomes (black circles) represented, along with average sequence composition of genes (green), DNA transposons (purple) and RNA transposons (blue). The relative positions of DSBs (red stars) in wild type euchromatin and heterochromatin, and in *kyp suvh5 suvh6* heterochromatin, are indicated.

We tested the role of non-CG DNA methylation and H3K9me2 in REC8 loading using *kss* mutants (Stroud et al. 2013, 2014). We observed remodeling of REC8 occupancy in *kss*, with regions that lose H3K9me2 and non-CG DNA methylation showing cohesin depletion and gain of SPO11-1-oligos. This is consistent with cohesin and heterochromatin jointly suppressing meiotic DSBs, and is reflected by the shared preference of REC8 and nucleosomes for GC-rich sequences. In contrast, SPO11-1-oligo and crossover hotspots are AT-biased. Unlike in fission yeast *clr4* or *swi6* mutants, which lose heterochromatic H3K9me2 and centromere cohesion (Bernard et al. 2001; Nonaka et al. 2002; Mizuguchi et al. 2014; Ellermeier et al. 2010), Arabidopsis *kss* mutants retain sufficient centromeric REC8 to maintain sister cohesion. Hence, Arabidopsis more closely resembles mouse *suv39h1 suv39h2* H3K9me2 mutants, where mitotic cohesin is recruited to heterochromatin, but with remodeling on major versus minor satellite repeats (Koch et al. 2008; Guenatri et al. 2004). We propose that in plant and mammalian genomes additional heterochromatic features act redundantly with H3K9me2 to recruit and maintain cohesin in the centromeres. Interestingly, we observed SPO11-1 ChIP enrichment in the centromeres, coincident with elevated REC8 and nucleosome density. Hence, although SPO11-1 is axis-enriched in proximity to the heterochromatic loops, these sequences are not efficiently recruited to form DSBs or crossovers during meiosis (Fig. 7).

Transcription drives cohesin occupancy in diverse eukaryotes (Lengronne et al. 2004; Bausch et al. 2007; Sun et al. 2015; Busslinger et al. 2017). Consistently, we observed a negative relationship between transcription level and REC8-cohesin occupancy within Arabidopsis genes. Hi-C contact maps in fission and budding yeast, which like plants lack CTCF, display contact associations at gene scale termed ‘globules’ and ‘crinkles’ that relate to transcription (Hsieh et al. 2015; Mizuguchi et al. 2014). High-resolution Hi-C studies in Arabidopsis have revealed contact associations over single genes, where 5' and 3' ends interact (Liu et al. 2016). The role of cohesin in intragenic contacts is unknown, although this may relate to the REC8 enrichment we observed within gene bodies. We identified transcriptionally upregulated genes and transposons in *kss*, which showed depletion of REC8 and gain of SPO11-1-oligos. This is consistent with an important role for transcription in shaping cohesin occupancy in plants. It is also important to consider that plant heterochromatin is actively transcribed by Pol IV and Pol V RNA polymerases, which produce short transcripts required for RNA-directed DNA methylation (Law and Jacobsen 2010). Hence, it will be interesting to explore the effects of heterochromatic transcription on cohesin occupancy in plant centromeric regions.

Recombination suppression in proximity to the centromeres by cohesin and heterochromatin is important for fertility, as crossovers within these regions can cause aneuploidy (Lamb et al. 2005; Rockmill et al. 2006). Plant centromeres are flanked by large domains of transposon-dense pericentromeric heterochromatin (Law and Jacobsen 2010). As a consequence, plant chromosomes show pronounced telomere–centromere gradients of recombination, epigenetic modifications, cohesin enrichment, compartment identity and gene/transposon composition (Higgins et al. 2012; Feng et al. 2014; Choi et al. 2018). These gradients likely exert a profound effect on sequence diversity along the length of plant chromosomes and contribute to their functional stratification and evolution. Indeed, genetic variation in *REC8* and meiotic axis genes has been identified as targets of selection during polyploid evolution in *Arabidopsis arenosa* (Yant et al. 2013; Wright et al. 2015). This effect is likely via modification of crossover patterns and stabilization of polyploid chromosome inheritance (Yant et al. 2013; Wright et al. 2015). Hence, far from being static components of plant genome architecture, REC8-cohesin and the meiotic axis dynamically evolve and influence patterns of recombination and diversity.

## Acknowledgements

We thank Mathilde Grelon for the MLH1 antibody, Paul Fransz for discussions concerning cytological data, Chang Liu for providing Hi-C data and the Gurdon Institute for access to microscopes. Research was supported by grants from the ERC (SynthHotSpot) and BBSRC (BB/L006847/1 and BB/N007557/1), an EMBO long term postdoctoral fellowship (ALTF 807-2009) and a Gatsby Foundation Sainsbury studentship GAT3401. The authors declare no competing interests.

## Author Contributions

Conceptualization: CL, AJT, KC, JH, FCHF, IRH. Software: AJT, XZ, IRH. Investigation: CL, KC, SDT, PCK, ARB, JH. Writing: CL, AJT, KC, ARB, KO, JH, FCHF, IRH.

